# Hydration and H/D exchange-dependent infrared signatures of the GCN4 leucine zipper

**DOI:** 10.64898/2026.06.26.731617

**Authors:** Hinduja Bhuvanendran, Carla M. Brunner, Hannes Kempf, Jonathan L. Moro, Eugenio Roubieu, Florian Turbant, André Mateus, Haifeng Lin, Lakshmi Das, Dmitry Malyshev, Ben Johns, Antonietta Parracino, Annalisa Pastore, Judith Peters, Aitziber L. Cortajarena, Laura Zanetti-Polzi, Nicolò Maccaferri

## Abstract

Attenuated total reflectance Fourier-transform infrared (ATR-FTIR) spectroscopy of proteins in aqueous solution is often limited by water absorption and other optical artifacts. To overcome these limitations, we evaluated the structural features and hydrogen-deuterium exchange (HDX) kinetics of the *α*-helical protein GCN4 in both hydrated (wet) and vacuum-dried (dry) states. While solvent heavily mask the second-derivative spectra of wet samples, vacuum drying yielded a thin, protein-rich film on the ATR crystal, significantly enhancing the signal-to-noise ratio and resolving the protein features without altering the native structure. Dry-state analysis clearly resolved the Amide I, Amide II, and deuterium-shifted Amide II’ (1450 cm^−1^) bands. Notably, second-derivative analysis of the dry spectra of the HDX samples revealed a bimodal Amide I distribution consisting of a stationary band at 1653 cm^−1^ from the solvent-inaccessible regions and an isotopically sensitive band shifting from 1648 cm^−1^ to 1644 cm^−1^ from solvent-accessible regions. These results demonstrate that vacuum-dried ATR-FTIR spectroscopy effectively eliminates solvent masking, providing the spectral clarity required to resolve discrete *α*-helical sub-populations after deuteration.

## Introduction

Understanding protein conformational stability in different solvent environments is essential for linking structure to function, as hydration strongly influences hydrogen-bonding networks, electrostatic interactions, and backbone flexibility [1–7]. These effects are most directly reflected in the protein secondary structure, which is highly sensitive to environmental perturbations such as solvent composition and isotopic substitution [5, 8–11]. Among the available structural techniques, attenuated total reflection Fourier transform infrared (ATR-FTIR) spectroscopy has become a widely used and powerful method for probing protein secondary structure in both solution and film states [12–14].

ATR-FTIR is particularly valuable because the vibrational modes of the protein backbone are highly sensitive to local structural organization. In particular, the Amide I ( ∼1649–1652 cm^−1^) band (dominated by C=O stretching) is strongly correlated with secondary structural motifs such as *α*-helices, *β*-sheets, and random coils, making it one of the most widely used spectroscopic markers for protein structure determination [9]. The Amide II ( ∼1543–1547 cm^−1^) band (NH bending coupled with CN stretching) provides complementary information on backbone conformation and hydrogen-bonding environments [9, 10]. Its sensitivity to hydrogen–deuterium exchange also makes it a useful probe for monitoring solvent accessibility and protein structure [15, 16]. Because of this combination of structural sensitivity and experimental accessibility, ATR-FTIR is widely used to study protein folding, aggregation, and solvent-induced conformational changes under near-physiological conditions [17, 18].

A major challenge in ATR-FTIR studies of proteins in aqueous environments is the strong absorption of liquid water in the mid-infrared region, which overlaps significantly with both Amide I and Amide II bands [19, 20]. This complicates baseline correction and can affect the reliable assignment of secondary structure related the measured spectral features. To overcome this limitation, isotopic substitution with D_2_O is commonly used to shift solvent- and protein-associated vibrational frequencies and reduce spectral overlap [15, 16].

A particularly effective approach is partial H_2_O/D_2_O exchange, which has emerged as a practical strategy to improve spectral resolution without requiring complete deuteration [11, 21, 22]. Even intermediate levels of hydrogen–deuterium exchange are sufficient to shift the Amide II band toward the lower-frequency Amide II^*′*^ region, thus reducing overlap with the Amide I envelope [11, 15, 16]. This partial separation improves the resolution of protein-derived spectral components and enables more reliable deconvolution of overlapping bands. As a result, secondary structure estimation, particularly the quantification of *α*-helical and *β*-sheet content, becomes more robust and less sensitive to solvent-induced distortions, baseline uncertainties, and spectral congestion. Consequently, partial deuteration is widely adopted in ATR-FTIR studies of proteins in complex aqueous environments [11]. This study focuses on GCN4 as a biologically and structurally important model system. GCN4 is a yeast transcription factor that plays a central role in regulating amino acid biosynthesis and mediating cellular responses to nutrient starvation [23–26]. It acts as a master regulator of metabolic adaptation by activating genes involved in amino acid and purine metabolism, thereby coordinating the general amino acid control response [27, 28]. Structurally, GCN4 contains a homodimeric leucine zipper domain characterized by a highly basic DNA binding portion (basic region) and ending in a coiled-coil folding motif in which C-terminal *α*-helices associate through hydrophobic leucine residues and interhelical packing interactions to form a stable functional complex (Fig. 1a) [29, 30].

**Figure 1:**
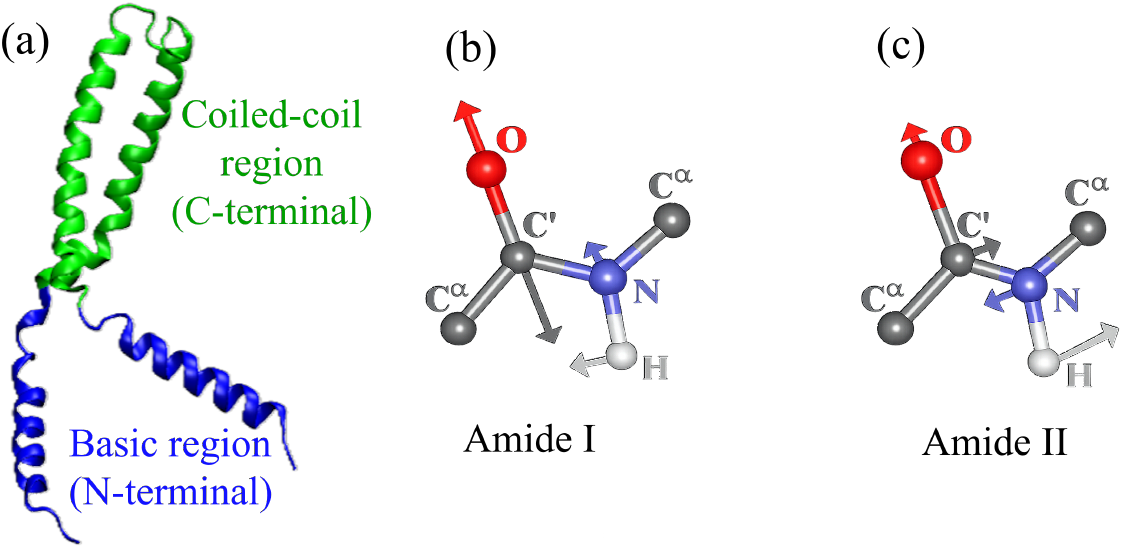
Structural and vibrational features of GCN4 and protein backbone modes. (a) Schematic representation of the leucine zipper domain of transcription factor GCN4 showing the N-terminal basic region (blue) responsible for sequence-specific DNA binding and the C-terminal coiled-coil domain that mediates dimerization and stabilizes the functional dimer–DNA complex. (b) Schematic of the amide I vibrational mode in a protein bond, primarily arising from C=O stretching of the backbone. (c) Schematic of the amide II vibrational mode, dominated by N–H in-plane bending coupled with C–N stretching. Directional arrows indicate the principal atomic displacement vectors associated with each vibrational mode.

The leucine zipper domain of GCN4 has served as a model in many studies aimed at elucidating the principles underlying coiled-coil structure and folding [31]. Importantly, GCN4 is not only a structural benchmark but also a prototypical representative of the AP-1 transcription factor family in humans, including proteins such as c-Fos and c-Jun [32, 33]. These proteins share the same leucine zipper dimerization architecture and rely on coiled-coil formation for DNA binding and transcriptional regulation. As such, insights obtained from GCN4 are broadly transferable to AP-1 family proteins, particularly in understanding dimer selection, stability, and environment-dependent conformational regulation. This makes GCN4 a valuable minimal model for studying biologically relevant coiled-coil transcription factors [32].

GCN4 is also an excellent system for vibrational spectroscopy because of its high *α*-helical content and well-defined structure, which produces strong and interpretable Amide I signatures directly reporting on folding, dimerization, and environmental perturbations. Despite extensive structural characterization by crystallography [34, 35], NMR [36, 37], and computational approaches [38], the influence of hydration and hydrogen–deuterium exchange on its infrared vibrational response remains still partially understood.

Addressing this gap is important not only for improving spectral interpretation in coiled-coil systems but also for strengthening the reliability of ATR-FTIR-based secondary structure analysis under physiologically relevant solvent conditions.

In this work, we systematically investigate the effects of hydration and isotopic substitution on the ATR-FTIR spectra of the GCN4 leucine zipper using a combination of H_2_O and H_2_O/D_2_O solvent systems. Measurements were first performed under hydrated (wet) conditions to assess the impact of hydrogen-deuterium exchange (HDX) on the Amide I and Amide II spectral signatures. The samples were subsequently analyzed in a vacuum-dried (dry) state to minimize solvent interference and enable direct observation of isotope-dependent changes in protein backbone vibrations, including the Amide II to Amide II^*′*^ transition. This experimental design allows solvent-derived spectral contributions to be distinguished from intrinsic protein features and reduces the risk of misinterpreting solvent artifacts as structural changes. Finally, second-derivative analysis of the ATR-FTIR spectra in the Amide region was employed to compare hydrated and dried samples across H_2_O and H_2_O/D_2_O conditions, providing insights into HDX behavior, solvent accessibility, and the preservation of the underlying *α*-helical coiled-coil structure. Together, these measurements establish a framework for interpreting ATR-FTIR spectra of coiled-coil proteins and demonstrate how controlled isotopic substitution can reveal structural heterogeneity that remains unresolved in H_2_O alone.

## Materials and Methods

### Protein Expression and Purification

To prepare the GCN4 bZIP samples from *Saccharomyces cerevisiae*, comprising the DNA-binding region and leucine zipper domain, we expressed and purified the protein using the untagged pGCNK58 plasmid [39]. Competent *Escherichia coli* BL21 (DE3) cells were transformed with the plasmid and grown in Luria Broth at 37 °C. When the culture reached an optical density at 600 nm of approximately 0.5, protein expression was induced with 1 mM IPTG for 4 h. Then the cells were harvested by centrifugation and lysed via sonication in a buffer containing 25 mM HEPES (pH 7.5) and 50 mM NaCl. The lysate was clarified and purified using a 5 mL HiTrap SP FF ion-exchange column on an ÄKTA Go system, eluting with a linear NaCl gradient from 50 mM to 1 M. Fractions containing GCN4 were identified by SDS-PAGE, pooled, and concentrated to 1.5 mM. Finally, the protein was dialyzed against 25 mM HEPES, 20 mM NaCl and (pH 8.0) prior to downstream applications.

### Sample Preparation

The GCN4 bZIP stock solution (1.5 mM in 25 mM HEPES, 20 mM NaCl, H_2_O) and a corresponding protein-free buffer were utilized to prepare two parallel dilution series with H_2_O and D_2_O at room temperature (approximately 22 °C). To reach a maximum working concentration of 750 *µ*M, the stock was initially diluted 1:1 with both H_2_O and D_2_O. Subsequently, a serial dilution was performed using the respective solvent to generate a GCN4 concentration gradient of 750, 375, 187, 93, and 46 *µ*M. Each dilution was homogenized via repeated trituration (pipetting) to ensure solvent and solute uniformity.

The same protocol was used for diluting the buffer components; HEPES concentrations ranged from 12.5 mM to 0.78 mM, while NaCl concentrations ranged from 10 mM to 0.63 mM. With the above set of protein concentrations, we perform two sets of measurements: in H_2_O dilution conditions and in D_2_O dilution conditions. For H_2_O dilution conditions, all the concentration series had almost 99% H_2_O and the protein concentration is decreased with dilution. For D_2_O dilution conditions, the highest protein concentration has almost 50-50 H_2_O-D_2_O ratio and with dilution D_2_O percentage increases and H_2_O percentage decreases. The percentage of each component in D_2_O dilution conditions is reported in Table 1.

**Table 1:** Mole percentage (mol%) composition of GCN4 samples for measurements in D_2_O dilution conditions.

| GCN4 stock | GCN4 (mol%) | HEPES (mol%) | NaCl (mol%) | H<sub>2</sub>O (mol%) | D<sub>2</sub>O (mol%) |
| --- | --- | --- | --- | --- | --- |
| 750 μM | 0.0014 | 0.023 | 0.046 | 49.26 | 50.67 |
| 375 μM | 0.0007 | 0.011 | 0.023 | 24.63 | 75.34 |
| 187 μM | 0.0003 | 0.006 | 0.011 | 12.31 | 87.67 |
| 93 μM | 0.0002 | 0.003 | 0.006 | 6.16 | 93.83 |
| 46 μM | 0.0001 | 0.001 | 0.003 | 3.08 | 96.92 |

All protein samples and corresponding buffer controls were incubated for 2 h at room temperature to allow hydrogen–deuterium exchange of the amide backbone. ATR-FTIR measurements were performed immediately after the incubation period, with all samples maintained under identical ambient conditions.

### FTIR Measurements

ATR-FTIR spectra were acquired using a Bruker Vertex 80 spectrometer equipped with a diamond ATR crystal. Spectra were recorded over a spectral range of 400–4000 cm^−1^ at a resolution of 1 cm^−1^. For each measurement, 100 scans were recorded and Fourier-transformed to improve the signal-to-noise ratio.

Experimental measurements were conducted on a single sample aliquot using a dual-state (wet and dry) protocol, enabling direct comparison between the wet and dry states from the same sample. A 2 *µ*L volume of either the protein solution or the corresponding buffer was drop-cast onto the ATR crystal. Hydrated (wet) spectra were initially recorded; during this phase, the sample was tightly sealed with a specialized lid to prevent solvent evaporation and maintain a constant solute concentration. Following the wet measurement, the same aliquot was dehydrated *in situ* via vacuum-assisted evaporation. Dehydration was maintained until a pressure of below 5 hPa was achieved, ensuring the formation of a uniform, dry protein film for subsequent acquisition of dehydrated (dry) spectra.

To minimize potential carryover effects and cross-contamination, the concentration series was measured progressively from the lowest (46 *µ*M) to the highest (750 *µ*M). Between each sample, the ATR crystal was cleaned with deionized water and ethanol. To ensure a stable baseline and atmospheric transparency, a background spectrum was recorded under vacuum conditions prior to measurements. This procedure facilitated the removal of residual CO_2_ and water vapour contributions to the spectra, even though it was not possible to fully eliminate these peaks.

Finally, the corresponding buffer reference spectra, acquired under identical wet and dry states under vacuum conditions, were subtracted from the protein spectra to isolate the protein’s vibrational signatures for analysis. To improve visualization of the second-derivative spectra, the data were interpolated onto a finer wavenumber grid using cubic spline interpolation (interp1, MATLAB). This procedure generates a smooth, continuous representation of the spectra by fitting piecewise cubic polynomials between adjacent data points while maintaining continuity of the first and second derivatives. The interpolation was applied solely for visualization purposes and did not alter the underlying spectral data used for analysis.

## Results and Discussion

ATR-FTIR measurements were first performed on GCN4 samples diluted exclusively in H_2_O in order to establish the intrinsic vibrational profile of the protein under standard aqueous conditions. Spectra were collected in both wet and dry states to evaluate the influence of solvent removal on spectral quality and protein structure.

For wet H_2_O measurements, the dominant features arise primarily from water O–H vibrational modes as shown in Figures 2a and 2b. Subtraction of the buffer spectrum from the corresponding sample yielded spectra with noticeable but weak protein-related features, as shown in Figure 2c.

**Figure 2:**
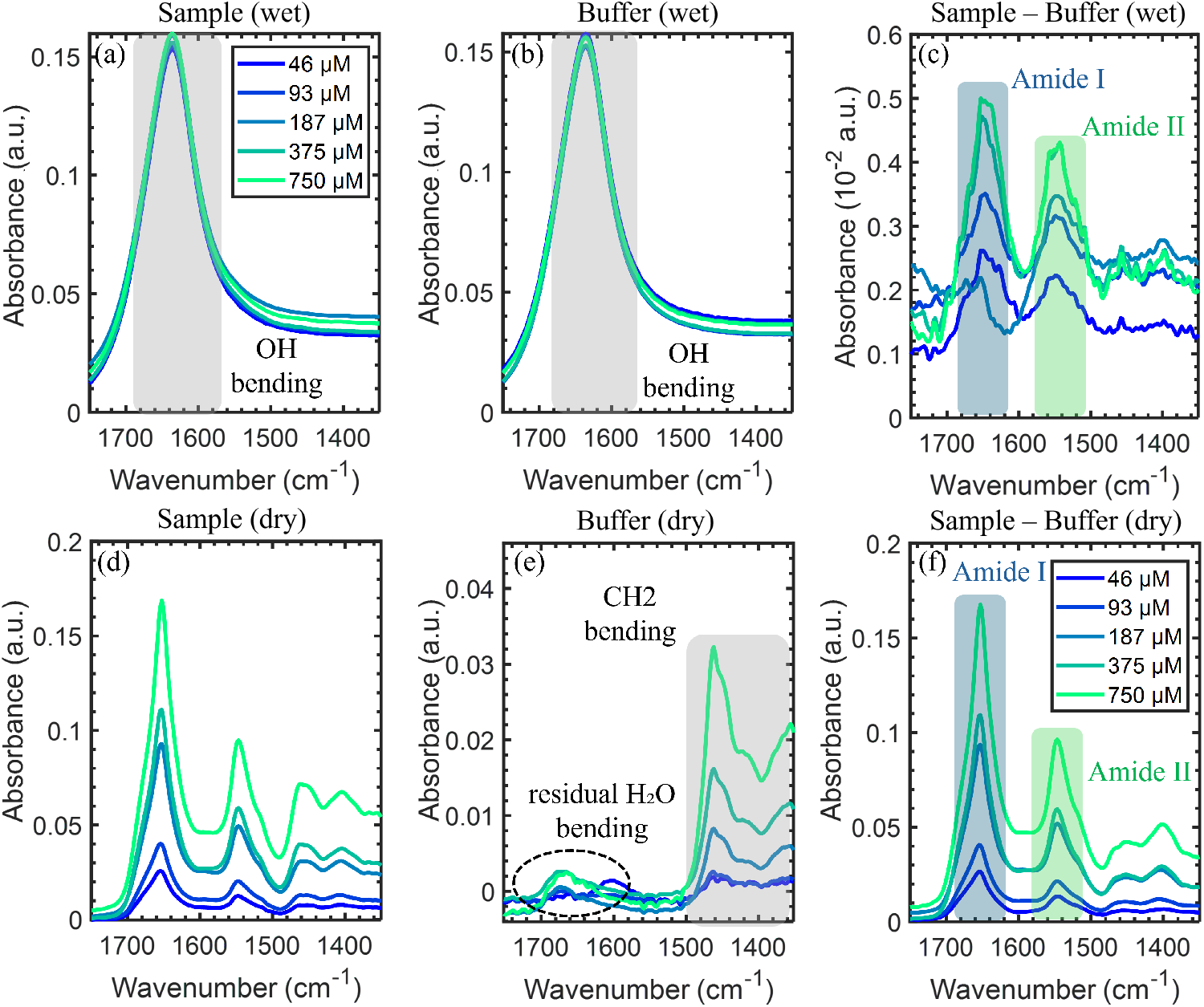
ATR-FTIR spectra of GCN4 in wet and dry states in H_2_O at different concentrations. Raw spectra of the (a) Sample (GCN4, HEPES and NaCl) and (b) buffer alone ( HEPES and NaCl) in the wet state, both dominated by the strong OH bending vibration of H_2_O. (c) Difference spectrum (sample minus buffer) in the wet state, resolving the characteristic Amide I and Amide II bands of the GCN4. Raw spectra of the (d) GCN4 sample and (e) buffer alone in the dry state, showing distinct spectral signatures from both GCN4 and the buffer components. (f) Dry-state difference spectrum (sample minus buffer), displaying the Amide I and Amide II bands with a significantly enhanced signal-to-noise ratio (SNR) relative to the wet-state difference spectrum 2c.

The Amide I band, primarily associated with protein backbone C=O stretching vibrations, is observed as a broad peak from ≈ 1653 cm^−1^ to 1639 cm^−1^ in the hydrated sample as shown in Figure 2c. The Amide II band, mainly originating from coupled NH bending and CN stretching modes, also shows a broad peak from ≈1559 cm^−1^ to 1543 cm^−1^. Although these features are detectable in the wet spectra, the overall signal intensity remained low because of strong overlapping absorption from bulk water, particularly the OH bending mode centered near ∼1630 cm^−1^.

ATR-FTIR spectra acquired from a protein solution under vacuum with the ATR lid closed are dominated by strong water absorption, resulting in a weak protein signal and a reduced signal-to-noise ratio (SNR). In attenuated total reflectance (ATR) spectroscopy, the infrared beam undergoes total internal reflection within the ATR crystal, generating an evanescent wave that penetrates only a short distance into the sample, typically on the order of micrometers. When a hydrated liquid layer is present on the ATR surface, the evanescent field primarily probes water molecules, whose broad and intense absorption bands can obscure weaker protein vibrational modes. During vacuum drying with an open or effectively ventilated configuration, water is progressively removed from the penetration depth region, leading to the formation of a thin protein-rich film directly in contact with the crystal. As a result, a larger fraction of the evanescent field interacts with protein rather than water, minimizing spectral masking effects and enhancing the relative intensity of protein absorption bands. Consequently, dry-state ATR-FTIR measurements exhibit significantly improved SNR and clearer protein spectral features than those obtained under hydrated conditions.

So, to improve signal to noise ratio and reduce solvent interference, the same samples were analyzed after vacuum drying. Removal of bulk water significantly reduced the background absorption and improved the signal-to-noise ratio across the amide region. In the dry-state spectra, Figures 2d and 2e show the sample and buffer measurements, respectively, and in the buffer spectrum, a residual tail of water absorption was still observable in the 1600–1700 cm^−1^ region. Figure 2f presents the corresponding buffer-subtracted spectrum and displayed well-resolved vibrational bands with substantially greater intensity than the corresponding wet measurements. The Amide I band was centered at 1653 cm^−1^, while the Amide II band appeared at 1547 cm^−1^. Compared with the dry state, hydration induced pronounced broadening of both bands due to hydrogen-bonding interactions with surrounding water molecules. In contrast, in the dry state these interactions are largely removed, resulting in sharper peaks converging toward a single dominant vibrational mode.

Importantly, no additional structural bands or major spectral distortions were observed after dehydration, indicating that vacuum drying did not significantly perturb the dominant structure of GCN4. The similarity between the wet and dry spectra demonstrates that the protein retains its characteristic spectral signature despite solvent removal, while the dry-state measurements provide substantially improved spectral clarity for subsequent analyses.

### ATR-FTIR Analysis of GCN4 in Mixed H_2_O/D_2_O Solvents and Spectral Overlap Effects

To investigate hydrogen–deuterium exchange in GCN4, the protein was progressively diluted using D_2_O and analyzed by ATR-FTIR spectroscopy under both wet and dry conditions. The amide II region is used as the primary probe for isotopic exchange because this vibration is strongly dependent on NH bending. Replacement of hydrogen with deuterium shifts this mode to lower frequencies, producing the characteristic Amide II’ band near ∼ 1450 cm^−1^ [9, 40].

As H_2_O and D_2_O are mixed during hydrogen-deuterium exchange (HDX), measurements taken in solution (wet conditions) are heavily dominated by solvent features. This spectral dominance is expected due to the solvent’s high molar concentration relative to the protein. Consequently, the raw spectra exhibit intense solvent interference, characterized by the H_2_O (OH) bending mode at 1630 cm^−1^ and the HOD bending mode in the 1450cm^−1^ region, as demonstrated in Figures 3a and 3b. Subtraction of the buffer spectrum from the corresponding sample yields weak protein-related features, as shown in Figure 3c. But the resulting protein spectrum exhibits very low absorbance intensity, approaching the experimental noise floor (∼ 10^−3^ absorbance units). Also, subtraction artifacts dominate the Amide II’ region, where nearly identical raw intensities between sample and buffer result in weak and unreliable residual signals. In mixed isotopic solvents, rapid proton exchange between H_2_O and D_2_O generates HOD, producing broad absorptions that significantly complicate background subtraction and obscure weak protein signals, especially in the encircled region in Figure 3c. The weak Amide I region is a broad peak from 1653 cm^−1^ to 1637 cm^−1^ and Amide II and Amide II’ is not clearly visible in the wet spectra due to the above mentioned reasons. At these concentrations, the sample and buffer spectra are nearly indistinguishable, indicating that the observed absorbance is dominated by solvent rather than protein backbone contributions.

**Figure 3:**
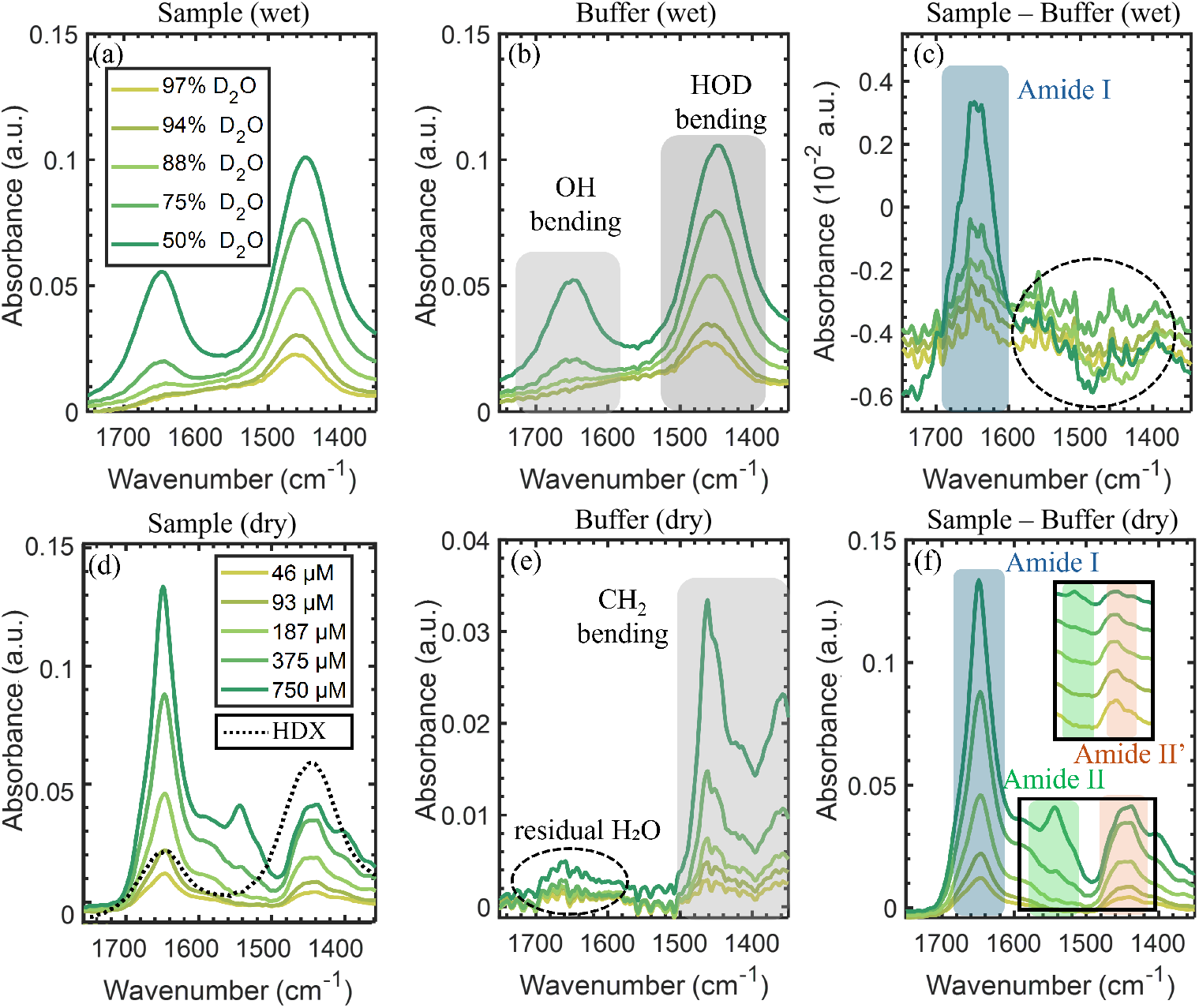
ATR-FTIR spectra of GCN4 in wet and dry states in different ratios of H_2_O:D_2_O at different concentrations of GCN4. Raw spectra of the (a) Sample (including GCN4, HEPES and NaCl) and (b) buffer alone ( HEPES and NaCl) in the wet state, both dominated by the strong OH bending vibration of H_2_O and HOD from H_2_O-D_2_O mixture. (c) Difference spectrum (sample minus buffer) in the wet state, resolving the characteristic Amide I of the GCN4. Amide II could not be resolved due to the high strength of HOD in this region. (d) Raw spectra of the sample with H_2_O*/*D_2_O mixture in dashed line to show protein and solvent peaks are at similar wavenumbers. (e) Raw spectra of buffer alone in the dry state, showing distinct spectral signatures of buffer. (f) Dry-state difference spectrum (sample minus buffer). The main plot displays the Amide I (blue shaded), Amide II (N–H bond, green shaded), and Amide II’ (N–D bond, orange shaded) bands. The inset details the normalized Amide II and II’ regions; both bands exhibit comparable intensities at approximately 50% D_2_O, and with increasing D_2_O, the Amide II band disappears, and the Amide II’ feature becomes dominant as at 97 % D_2_O .

We systematically compared spectra collected under dry conditions obtained after drying with the wet spectra. In the dry state, removal of bulk solvent substantially reduced overlapping OH and HOD absorptions, thereby improving the reliability of the protein spectra. The relative intensity between Amide II and Amide II’ is decreasing as more dilution with D_2_O, Amide II disappears, and Amide II’ is more prominent. In Figure 3d, the dotted black line shows the H_2_O-D_2_O mixture, highlighting a potential source of peak misinterpretation. As shown, the H_2_O-D_2_O mix spectrum, which does not contain any Amide I or Amide II contributions, nonetheless exhibits apparent features in the same spectral regions. These features arise from solvent OH and HOD bending vibrations, which overlap with the expected positions of the Amide I and Amide II’ bands.

The weak feature observed near ∼ 1670 cm^−1^ in the dry buffer spectrum (Figure 3e) arises from the high-frequency tail of the OH bending mode of residual water, which is broad and strongly dependent on the local hydrogen-bonding environment [41, 42].

Although weak residual solvent contributions remained visible, the overall spectral background is significantly lower than in wet measurements. After subtracting the buffer contribution, the resulting dry-state spectra revealed well-resolved protein vibrational features, including the Amide I band at 1649 cm^−1^ [11], the Amide II band at 1545 cm^−1^, and a distinct Amide II’ band centered near 1452 cm^−1^ as shown in Figure 3f. The main plot highlights the well-defined Amide I region alongside the distinct contributions from the Amide II (NH) and deuterated Amide II’ (ND) bands.To closely evaluate the isotope exchange efficiency, the inset focuses on the normalized Amide II and II’ regions. At 50% D_2_O, both bands exhibit comparable spectral intensities, reflecting a balanced mixture of protonated and deuterated states. As the DO content is systematically raised, the progressive replacement of amide protons by deuterons drives the total disappearance of the Amide II band, leaving the Amide II’ feature entirely dominant at 97% D_2_O.

As seen in H_2_O, wet and dry here as well, the wet is broader, and the dry has narrower, clearer peaks. The observed spectral evolution reflects the isotopic substitution of N-H groups by deuterium to form N-D, thereby increasing the reduced mass of the vibrating system and shifting the corresponding vibrational frequency to lower wavenumbers. These results also show a clear correlation between the Amide II to II’ ratio and the concentration of D_2_O.

### Secondary Structure Analysis from the Amide I Region

A common and reliable practice for a detailed examination of secondary structure is second-derivative analysis of the Amide I region. This technique effectively sharpens hidden peaks; note that peaks in conventional spectra will appear as dips in the second-derivative spectrum. Following buffer subtraction, this second-derivative processing resolves overlapping *α*-helical contributions that are otherwise obscured in conventional spectra.

As shown in Figure 4a, the Amide I band for the H_2_O dry state appeared as a single dominant feature across all concentrations. A zoomed view of the 1670 cm^−1^ region revealed a single dip at 1653 cm^−1^ as shown in Figure 4b, which is a characteristic feature of an *α*-helix; notably, the dashed box highlights a region where no other peaks are expected. In contrast, the wet spectra exhibited prominent solvent effects despite solvent and buffer subtraction. This interference is clearly visible within the dashed box in Figure 4c, a region that should ideally be free of spectral features. Monitoring this specific window serves as an effective method to verify whether observed peaks genuinely originate from the protein sample or are merely solvent artifacts. To investigate this further, the second-derivative peak positions from the wet measurements are compared with documented water vapour features in FTIR spectroscopy. The observed peaks closely matched characteristic vapour-associated positions at 1684, 1670, 1663, 1653, 1647, 1636, 1623, and 1616 cm^−1^ [43]. This strong correspondence supports assigning these sharp, concentration-independent features to water vapour rather than solvent- or protein-derived contributions. A standard approach to assess such interference is to examine the protein-free baseline region between 1700 and 1800 cm^−1^. While the wet spectra in this study exhibited several derivative features in this range—confirming residual solvent and water vapour contamination—the corresponding dry-state spectra remained relatively featureless. This clean baseline in the dry state demonstrates significantly reduced interference, confirming that the observed Amide I features genuinely originate from the protein.

**Figure 4:**
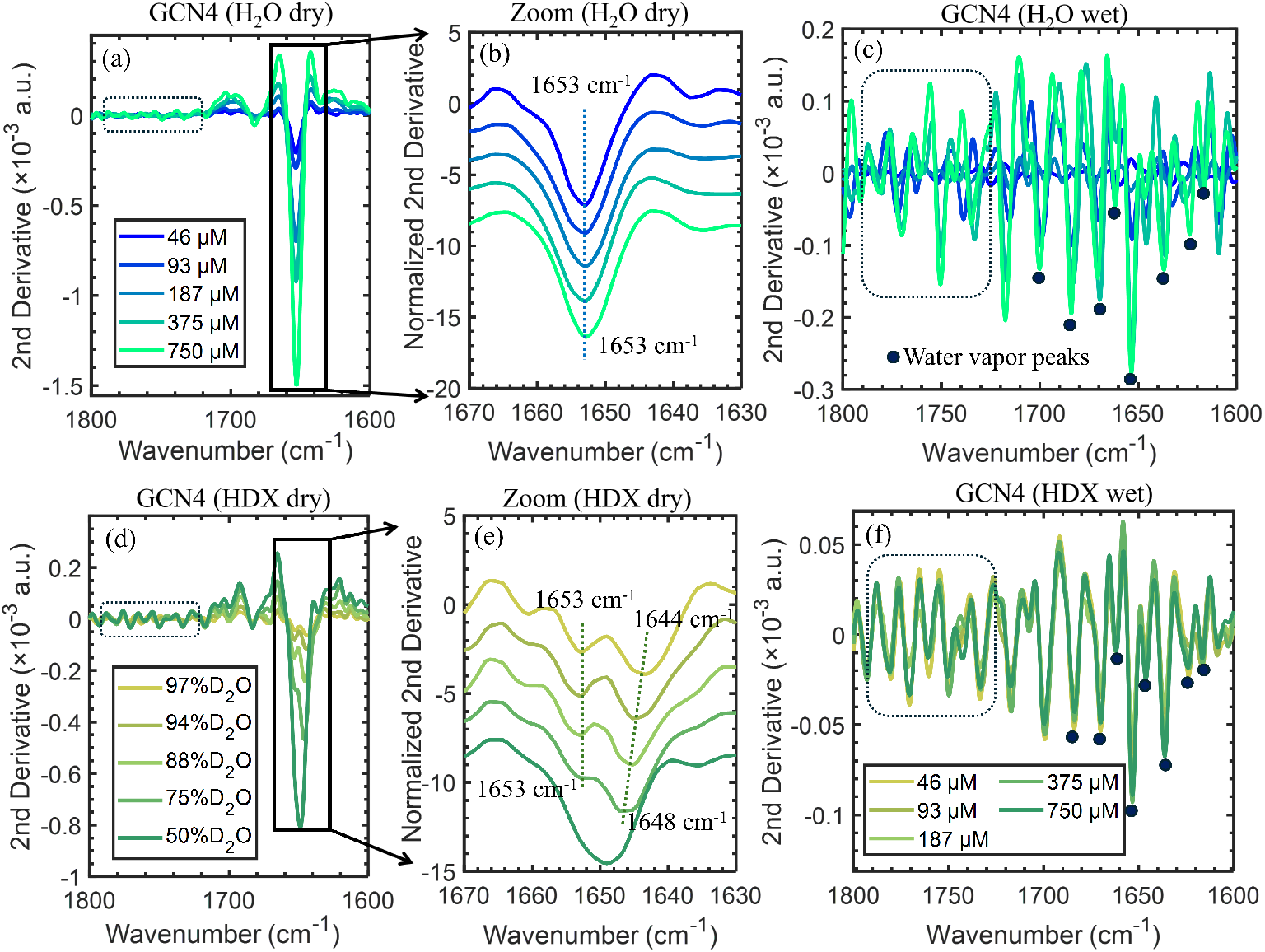
Second derivative spectra of the GCN4 protein at different concentrations prepared in H_2_O and H_2_O/D_2_O. (a) Second derivative spectra of GCN4 diluted in H_2_O (dry), which shows peaks at 1653 cm^−1^, and the dashed box shows no prominent peaks in the 1750-1800 cm^−1^ region. (b) Magnified and normalized view of the Amide I region (1670-1630 cm^−1^), highlighting changes in the protein secondary structure. (c) Second derivative spectra of GCN4 diluted in H_2_O (wet), which is mainly water vapour, and the dots show the water vapour peak positions mentioned in the text. (d) Second derivative spectra of GCN4 diluted in H_2_O*/*D_2_O (dry) and D_2_O percentage is 50%, 75%, 88%, 94%, and 97% as the concentration of the sample decreases. (e) Magnified and normalized view of the Amide I region (1670-1630 cm^−1^), highlighting changes in the protein secondary structure. A red shift (1648-1644 cm^−1^) in the peaks due to increased D_2_O content can be observed. (f) Second derivative spectra of GCN4 diluted in H_2_O*/*D_2_O (wet), which is mainly water vapour and the dashed box shows solvent features remain even after subtraction.

In contrast to the single dominant peak observed in the H_2_O dry state, the second-derivative peaks in the dry spectra of the HDX samples began to split at higher D_2_O ratios. This splitting is visible in Figure 4d and is more clearly resolved in the zoomed-in view of Figure 4e. While one peak remained stationary at 1653 cm^−1^ across the entire dilution series, a second peak initially located at 1648 cm^−1^ progressively red-shifted to 1644 cm^−1^ as the D_2_O content increased. The stationary feature at 1653 cm^−1^ can be assigned to the solvent-inaccessible core of the *α*-helix. Conversely, the shifting component 1648-1644 cm^−1^ can be attributed to the solvent-accessible regions of the *α*-helix, as its pronounced red-shift directly correlates with increasing D_2_O exposure. The relative intensities of the two bands evolve with both increasing D_2_O fraction and decreasing protein concentration. The lower-frequency component is intrinsically more intense under low D_2_O conditions, and therefore dominates the composite envelope at high protein concentration (750 *µ*M), resulting in an apparent band maximum closer to ∼ 1648 cm^−1^. With increasing D_2_O fraction, the intensities of the two components become comparable, enabling their spectral separation and more accurate resolution of the underlying bimodal distribution. This kind of behavior is typically interpreted as arising from the coexistence of distinct *α*-helical populations with different solvent accessibility[44]. Specifically, the invariant 1653 cm^−1^ band is compatible with solvent-inaccessible C=O groups and the isotopically sensitive 1648-1644 cm^−1^ band is compatible with solvent-accessible C=O groups. However, since Amide I frequencies are strongly modulated by hydrogen-bond geometry and strength, the above-mentioned features may not exclusively reflect solvent inaccessibility or accessibility.

In the measurements of HDX samples in wet conditions, the sharp second-derivative features are attributed neither to the solvent nor to the protein, but rather to residual atmospheric water vapour within the spectrometer’s optical path, as indicated by the dots in Figure 4f. As observed similarly in the H_2_O wet spectra (Figure 4c), the presence of features inside the dashed box confirms that these spectra are unsuitable for analyzing protein secondary structure. These sharp vapour absorptions dominate the second-derivative response, producing dips that are significantly stronger than any protein-related signal. Although weak Amide I features are visible in the raw spectra, they cannot be resolved in the second derivative because the protein band is broad with a low slope. Because second-derivative processing preferentially amplifies sharp, high-slope transitions, the resulting spectra are entirely dominated by atmospheric water-vapour artifacts.

## Conclusion

This study reports a comprehensive ATR-FTIR characterization of GCN4 across varying hydration states and isotopic solvent conditions, providing insight into both methodological influences and the protein’s intrinsic structural features.

Comparative analysis of wet and dry measurements in H_2_O shows that while the characteristic Amide I and Amide II) bands of GCN4 are consistently detectable, aqueous measurements are strongly affected by overlapping solvent absorption, particularly from water OH bending near ∼1630 cm^−1^. Vacuum drying significantly reduces this spectral interference, leading to improved signal-to-noise ratios and more clearly resolved vibrational features. Importantly, the close agreement between the wet and dry spectra indicates that dehydration does not induce a significant perturbation of the dominant secondary structure, confirming the protein’s robustness under the applied experimental conditions.

Experiments with H_2_O/D_2_O mixtures further highlight the challenges posed by solvent-driven spectral overlap. Rapid hydrogen-deuterium exchange generates HOD bending vibrations that introduce broad absorptions, substantially complicating buffer subtraction, particularly in the Amide I and Amide II regions. These effects reduce the reliability of wet-state difference spectra and limit sensitivity to isotopic exchange in dilute protein samples. In contrast, dry-state measurements effectively suppress solvent contributions, enabling clear observation of the Amide II → Amide II’ transition, which provides a direct spectroscopic marker of H/D exchange in the protein backbone. The Amide I band remains centered around 1653 cm^−1^ in H_2_O and H_2_O/D_2_O, with minor shifts observed upon isotopic substitution to 1649 cm^−1^. Although D_2_O substitution induces modest redshifts in Amide I and Amide II’ bands, these changes reflect isotopic mass effects and solvent accessibility rather than global unfolding or loss of secondary structure in the conventional spectra. Secondary structure analysis of the Amide I region confirms that GCN4 retains a predominantly *α*-helical coiled-coil architecture across all conditions studied. The HDX measurements indicate the coexistence of solvent-accessible and solvent-inaccessible *α*-helical populations. Increasing D_2_O content leads to a gradual redshift of solvent-exposed helices, whereas a structurally protected core remains centered at 1653 cm^−1^, consistent with a stable coiled-coil architecture that preserves its secondary structure under all conditions examined.

Overall, the results demonstrate that GCN4 maintains a stable *α*-helical coiled-coil framework under both hydration and partial deuteration, while its vibrational signatures are strongly modulated by solvent environment and exchange dynamics. Interestingly, the study shows that vacuum-dried ATR-FTIR provides a reliable and higher-resolution approach for monitoring protein backbone vibrations and hydrogen–deuterium exchange by minimizing solvent-induced optical artifacts without altering the underlying structural integrity of the protein.

## Acknowledgements

This research was supported by the European Innovation Council Pathfinder Open project “iSenseDNA” (Grant No. 101046920), Kempestiftelserna (Grant No. Grant JCK-3122) and the UCMR “Excellence by Choice”postdoctoral program at Umeå University (Grant No. Grant No. JCK-2130.3) funded by Kempestiftelserna. The authors thank the Vibrational Spectroscopy Core Facility at Department of Chemistry and Chemical Biological Centre, Umeå University, for instrument access and support, and András Gorzsás and Andreas Barth for fruitful discussions.

## References

(1) Bellissent-Funel, M.-C.; Hassanali, A.; Havenith, M.; Henchman, R.; Pohl, P.; Sterpone, F.; van der Spoel, D.; Xu, Y.; Garcia, A. E. Chemical Reviews 2016, 116, 7673–7697.

(2) Ball, P. Chem Rev 2008, 108, 74–108.

(3) Levy, Y.; Onuchic, J. N. Annu Rev Biophys Biomol Struct 2006, 35, 389–415.

(4) Chaplin, M. Nat Rev Mol Cell Biol 2006, 7, 861–866.

(5) Jackson, M.; Mantsch, H. H. Crit Rev Biochem Mol Biol 1995, 30, 95–120.

(6) Arrondo, J. L. R.; Muga, A.; Castresana, J.; Goñi, F. M. Progress in Biophysics and Molecular Biology 1993, 59, 23–56.

(7) Arrondo, J. L. R.; Gońi, F. M. Prog Biophys Mol Biol 1999, 72, 367–405.

(8) Prabhu, N.; Sharp, K. Chemical Reviews 2006, 106, 1616–1623.

(9) Barth, A. Biochim Biophys Acta 2007, 1767, 1073–1101.

(10) Kong, J.; Yu, S. Acta Biochim Biophys Sin 2007, 39, 549–559.

(11) De Meutter, J.; Goormaghtigh, E. European Biophysics Journal 2021, 50, 613–628.

(12) Luthra, S.; Kalonia, D. S.; Pikal, M. J. Journal of Pharmaceutical Sciences 2007, 96, 2910–2921.

(13) Tintor, Ð.; Ninković, K.; Milošević, J.; Polović, N. Ð. Vibrational Spectroscopy 2024, 134, 103726.

(14) Tatulian, S. A. In Lipid–Protein Interactions: Methods and Protocols; Methods in Molecular Biology, Vol. 974; Humana Press: Totowa, NJ, 2013, pp 177–218.

(15) Maréchal, Y., The Hydrogen Bond and the Water Molecule; Elsevier: Amsterdam, 2007.

(16) DeFlores, L. P.; Ganim, Z.; Nicodemus, R. A.; Tokmakoff, A. J Am Chem Soc 2009, 131, 3385–3391.

(17) Shivu, B.; Seshadri, S.; Li, J.; Oberg, K. A.; Uversky, V. N.; Fink, A. L. Biochemistry 2013, 52, 5176–5183.

(18) Miller, L. M.; Bourassa, M. W.; Smith, R. J. Biochimica et Biophysica Acta (BBA) – Biomembranes 2013, 1828, 2339–2346.

(19) López-Lorente, Á. I.; Mizaikoff, B. Analytical and Bioanalytical Chemistry 2016, 408, 2875–2889.

(20) Tatulian, S. A. Biochemistry 2003, 42, 11898–11907.

(21) Giubertoni, G.; Bonn, M.; Woutersen, S. The Journal of Physical Chemistry B 2023, 127, 8086–8094.

(22) Baello, B. I.; Pancoska, P.; Keiderling, T. A. Analytical Biochemistry 2000, 280, 46–57.

(23) Hinnebusch, A. G. Annual Review of Microbiology 2005, 59, 407–450.

(24) Natarajan, K.; Meyer, M. R.; Jackson, B. M.; Slade, D.; Roberts, C.; Hinnebusch, A. G.; Marton, M. J. Molecular and Cellular Biology 2001, 21, 4347–4368.

(25) Hinnebusch, A. G. Annual Review of Microbiology 2005, 59, 407–450.

(26) Natarajan, K.; Meyer, M. R.; Jackson, B. M.; Slade, D.; Roberts, C.; Hinnebusch, A. G. Molecular and Cellular Biology 2001, 21, 4347–4368.

(27) Selman, C.; Tullet, J. M. A.; Wieser, D.; Irvine, E.; Lingard, S. J.; Choudhury, A. I.; Claret, M.; Al-Qassab, H.; Carmignac, D., et al. Science 2009, 326, 140–144.

(28) Petti, A. A.; McIsaac, R. S.; Ho-Shing, O.; Bhatt, D.; Bhatt, D.; Bhatt, D. PLOS Genetics 2022, 18, e1010505.

(29) O’Shea, E. K.; Klemm, J. D.; Kim, P. S.; Alber, T. Science 1991, 254, 539–544.

(30) Stitzel, M. L.; Durso, R.; Reese, J. C. Genes & Development 2001, 15, 128–133.

(31) Lumb, K. J.; Carr, C. M.; Kim, P. S. Biochemistry 1994, 33, 7361–7367.

(32) Sellers, J. W.; Struhl, K. Nature 1989, 341, 74–76.

(33) Stitzel, M. L.; Durso, R.; Reese, J. C. Genes & Development 2001, 15, 128–133.

(34) Yeates, T. O.; Kent, S. B. H. Annual Review of Biophysics 2012, 41, 41–61.

(35) Su, X.-D.; Zhang, H.; Terwilliger, T. C.; Liljas, A.; Xiao, J.; Dong, Y. Protein & Cell 2014, 5, 122–153.

(36) Kay, L. E. Journal of Magnetic Resonance 2011, 213, 477–491.

(37) Kaptein, R.; Boelens, R.; Scheek, R. M.; van Gunsteren, W. F. Biochemistry 1988, 27, 5389–5395.

(38) Devaurs, D.; Antunes, D. A.; Borysik, A. J. Journal of the American Society for Mass Spectrometry 2022, 33, 215–237.

(39) Gill, S. C.; von Hippel, P. H. Anal Biochem 2016, 175, S1–S12.

(40) Ganim, Z.; Chung, H. S.; Smith, A. W.; Cheatum, C. M.; Tokmakoff, A. Acc Chem Res 2008, 41, 432–441.

(41) Seki, T.; Chiang, K.-Y.; Yu, C.-C.; Bonn, M. Journal of Physical Chemistry Letters 2020, 11, 8459–8469.

(42) Bridelli, M. G. In Fourier Transforms – High-Tech Application and Current Trends; IntechOpen: 2017, pp 191–213.

(43) Zou, Y.; Ma, G. International Journal of Molecular Sciences 2014, 15, 10018–10033.

(44) Imamura, H.; Isogai, Y.; Takekiyo, T.; Kato, M. Biochimica et Biophysica Acta (BBA) -Proteins and Proteomics 2010, 1804, 193–198.

